# The eIF4B RNA recognition motif promotes higher-order organization of the translation initiation machinery during stress granule assembly

**DOI:** 10.64898/2026.08.26.747272

**Authors:** Jessica Bolivar, Nicholaus L. DeCuzzi, Elijah Kofke, Masaaki Sokabe, Katherine Beglinger, John G Albeck, Christopher S. Fraser

## Abstract

Cells respond to environmental stress by rapidly remodeling translation and assembling stress granules (SGs), which are dynamic ribonucleoprotein condensates that contain untranslated mRNAs, translation initiation factors, and 40S ribosomal subunits. Although the translation initiation factor eIF4B has been implicated in SG biology, the contribution of its highly conserved RNA recognition motif (RRM) to SG assembly has remained unclear. Here, we developed a quantitative live-cell imaging framework that resolves distinct kinetic phases of SG assembly at single-cell resolution and combines these measurements with single-cell analysis of protein synthesis. Using this approach, we show that disruption of the eIF4B RRM delays SG nucleation, slows SG assembly, and reduces the number of SGs formed, while having little effect on mature SG size. Biochemical analyses revealed that the RRM mutant retained high-affinity binding to both RNA and the 40S ribosomal subunit and exhibited only a modest reduction in eIF4A helicase stimulation activity but displayed altered RNA engagement, consistent with impaired RNA-dependent organization of the translation initiation machinery. Coupling SG kinetics with single-cell measurements of protein synthesis further revealed that delayed SG nucleation is associated with reduced translational repression during oxidative stress. Together, our findings identify the conserved eIF4B RRM as a regulator of productive higher-order organization of the translation initiation machinery and establish a quantitative framework for investigating how SG assembly and translational remodeling are coordinated during cellular stress.

## Introduction

Cells respond to environmental stresses, including heat shock and oxidative stress, by rapidly remodeling protein synthesis and sequestering mRNAs into stress granules (SGs).^1^ SGs are non-membrane–bound ribonucleoprotein condensates composed of untranslated mRNAs, RNA-binding proteins, 40S ribosomal subunits, and eukaryotic initiation factors (eIFs).^2,3^ SGs form when stalled translation initiation complexes accumulate on mRNAs, giving rise to dynamic ribonucleoprotein condensates.^4^ RNA molecules recruited into condensates self-organize, generating an RNA shell stabilized by intermolecular RNA–RNA interactions.^5^ Through these dynamic interactions, SGs coordinate cellular adaptation to stress by regulating the translational machinery while influencing mRNA storage, remodeling, and fate.^2^ Given their central role in cellular response to stress, dysregulated SG assembly has been implicated in cancer, neurodegeneration, and viral infection, underscoring the importance of understanding the molecular mechanisms that govern SG assembly and dynamics.^6–8^

Translation initiation is intimately coupled to SG assembly.^4,9^ During the integrated stress response, phosphorylation of eIF2α inhibits ternary complex formation, leading to the accumulation of stalled translation initiation complexes that seed SG assembly.^3^ However, SGs can also form through eIF2α-independent mechanisms, indicating that multiple pathways converge to regulate their formation.^10^ Consistent with this idea, several translation initiation factors have emerged as important regulators of SG dynamics.^11–13^ For example, eIF4A functions as an ATP-dependent RNA chaperone that limits mRNA condensation and SG formation^5^, whereas disruption of additional components of the translation initiation machinery also promotes SG assembly.^10,14^ In parallel, RNA-binding proteins including TIA1, TIAR, ataxin-2, and G3BP1/2 contribute to SG nucleation, with G3BP1 serving as a central organizer whose overexpression induces SG-like structures whereas its depletion blocks SG assembly.^15–20^

The eIF4F complex cooperates with accessory proteins including eIF4B, eIF4H, and poly(A)-binding protein (PABP) to promote translation initiation.^21,22^ eIF4B enhances the duplex-unwinding activity of eIF4A and, together with eIF4G, converts eIF4A into a processive helicase that promotes translation of structured mRNAs.^23–27^ Despite the evolutionary conservation of its RNA recognition motif (RRM), however, its biological function has remained surprisingly elusive. Domain dissection of yeast eIF4B demonstrated that the RRM is largely dispensable for translation initiation and cell growth, suggesting that it serves a regulatory rather than essential biochemical function.^28^ Recent studies have begun to provide insight into this longstanding question by showing that the intrinsically disordered regions of human eIF4B promote dynamic self-association at micromolar protein concentrations.^29^ In contrast, our recent work demonstrates that eIF4B drives regulated RNA-dependent formation of nanometer-scale clusters of the translation initiation machinery at nanomolar concentrations.^30^ These RNA-protein clusters (RPCs) greatly enhance eIF4A helicase activity. Together, these findings suggest that eIF4B functions not only as a helicase cofactor but also as an organizer of higher-order translation initiation complexes. Whether these assembly properties contribute to the cellular functions of eIF4B remains unknown.

Here, we investigated whether the higher-order assembly properties of eIF4B contribute to SG assembly during cellular stress. Using quantitative live-cell imaging and a single-cell analysis pipeline, we measured the kinetics of SG formation in cells overexpressing wild-type or RNA recognition motif (RRM) mutant eIF4B. Disruption of the RRM markedly impaired SG nucleation and assembly kinetics and altered the normal translational response to oxidative stress. Biochemical analyses demonstrated that the RRM mutant retained much of the core biochemical activity of eIF4B, including 40S binding, while exhibiting altered RNA engagement and reduced helicase stimulation. Together, our findings suggest that eIF4B contributes to the dynamic organization of the translation initiation machinery, thereby facilitating efficient stress granule assembly during cellular stress.

## Results

### A quantitative single-cell framework reveals the kinetics of eIF4B-dependent stress granule assembly

To investigate the dynamics of eIF4B recruitment to SGs, we generated a stable HeLa cell line expressing N-terminally GFP-tagged wild-type eIF4B (WT-GFP-eIF4B) under the control of a tetracycline (TET)-inducible promoter. Following induction, WT-GFP-eIF4B was expressed at levels comparable to endogenous eIF4B (**Figure S1D**). Prior to stress, WT-GFP-eIF4B was diffusely distributed throughout the cytoplasm. Upon sodium arsenite treatment, the signal rapidly localized into punctate structures within 30 minutes (**Figure 1A**). To confirm SG localization, we assessed colocalization between WT-GFP-eIF4B and G3BP1, a well-established SG marker, via immunofluorescence at the end of the live-cell imaging time course (**Figure S1A**). The Mander’s overlap coefficient for WT-GFP-eIF4B and G3BP1 was 0.81, indicating robust colocalization and suggesting that both proteins are similarly sequestered into SGs (**Figure S1B**).

**Figure 1.**
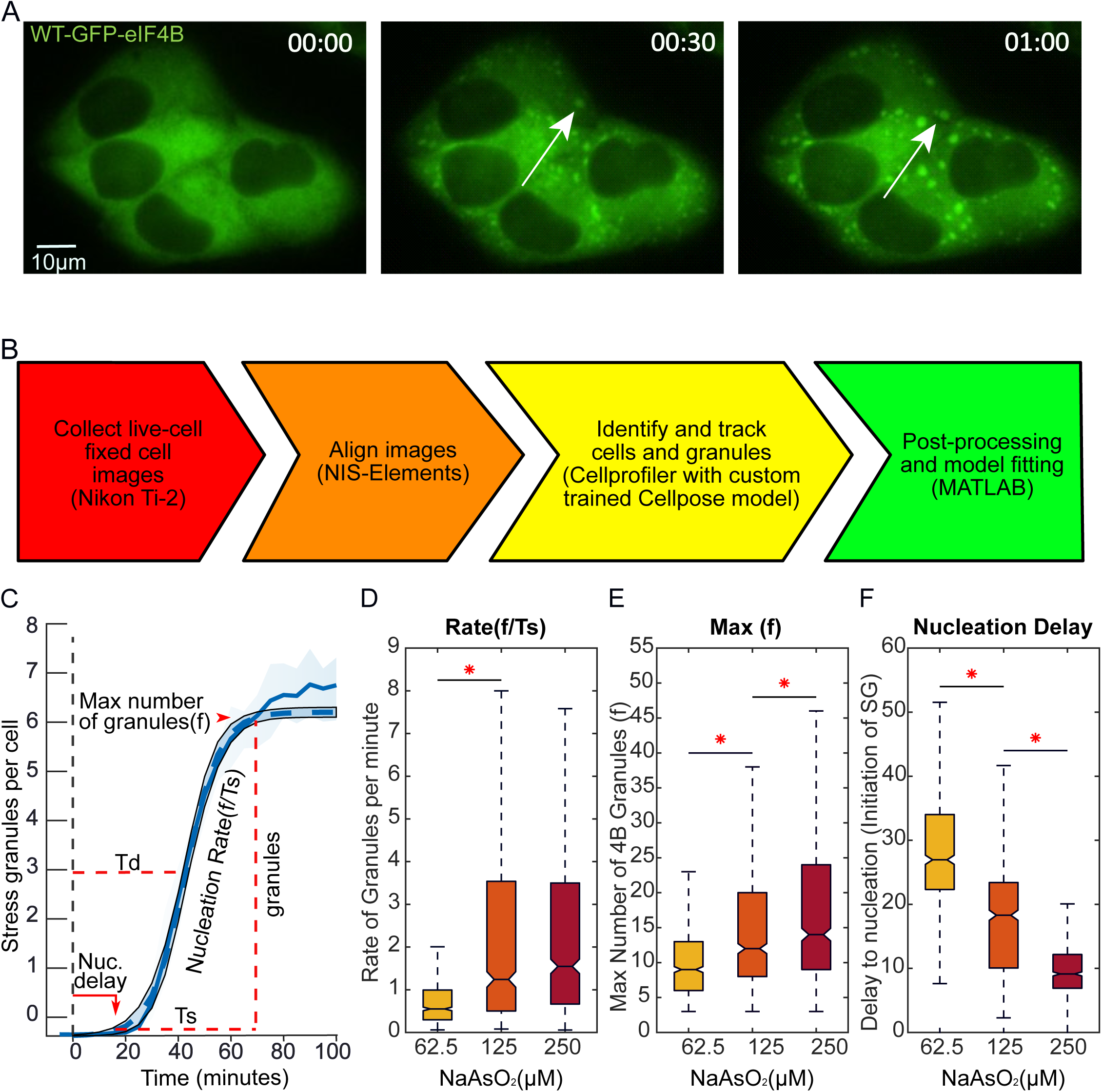
Quantitative single-cell analysis of GFP-eIF4B stress granule assembly. (A) Time-lapse imaging of stress granule formation in HeLa cells expressing WT-GFP-eIF4B following treatment with 125 μM sodium arsenite. The white arrow identifies an example of a SG formed during the time course (shown in hours). Scale bar is 10μm. (B) Workflow for quantitative analysis of SG assembly. Live-cell and immunofluorescent (fixed) images were acquired, shifts in the images are corrected, images are analyzed by a Cell Profiler pipeline that uses a custom Cellpose neural net trained to identify SGs; and the tracked SG data are imported into MATLAB where it is filtered for erroneous data and used to make plots and fit a model to determine the rate of SG assembly. (C) SG kinetics are extracted from live-cell data.^31^ This equation is fit to the single cell SG data to determine: Max number of SGs per cell (f), Rate of SG formation (f/Ts), and SG nucleation delay (Td – (1/2)Ts). Graph shows the average number of SGs per cell over time. The dark blue line is the mean of SGs and the shaded light blue area includes the 75 and 25 quantiles of mean. Dashed line is model fit to live-cell data with 95% confidence intervals. (D-F) Box and whisker plots showing single-cell WT-GFP-eIF4B rate, max number, and SG nucleation delay per cell. Colors indicate treatment with either 62.5 μM (Yellow), 125 μM (Orange), or 250 μM (Red) sodium arsenite (NaAsO_2_). Asterisk indicates P<0.005 between groups. Significance was determined via One-Way ANOVA comparison to compare the difference between each treatment group with a Dunnett multi-comparison protocol to account for false discovery rate. n>660 cells per condition pooled from 3 experimental replicates with at least 2 technical replicates each. See also Figure S1.

To quantitatively resolve SG assembly kinetics in individual cells, we developed a live-cell imaging pipeline that continuously tracked SG formation throughout the cellular stress response (**Figure 1B**). The analysis workflow integrated NIS Elements, CellProfiler, CellPose, and MATLAB to identify, quantify, and follow individual SGs over time. Granule formation kinetics were modeled using a modified version of an equation previously applied to describe caspase substrate cleavage in single cells.^31^

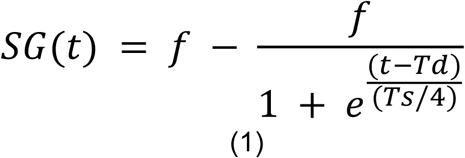

Fitting single-cell SG trajectories with this sigmoidal model, we were able to describe kinetic parameters of SG accumulation over time (**Figure 1C**), including the maximal number of SGs formed (**f**), the assembly time from first SG detection to maximal SG number (**Ts**), and the time from stress treatment to half-maximal SG formation (**Td**). From these parameters, we calculated the SG nucleation rate (**f/Ts**) and nucleation delay (**Td − Ts/2**), providing quantitative metrics for comparing SG assembly kinetics across conditions.

Using this quantitative framework, we next examined how the level of oxidative stress influences SG assembly in cells expressing WT-GFP-eIF4B. Individual SG trajectories were quantified following treatment with increasing concentrations of sodium arsenite (0–250 μM), and the kinetic model was fit to cells exposed to 62.5 μM, 125 μM, or 250 μM sodium arsenite.

Quantitative analysis revealed that increasing sodium arsenite concentrations accelerated SG assembly, increased the maximal number of SGs formed per cell, and shortened the SG nucleation delay (**Figure 1D–F**). The SG assembly rate (f/Ts) increased approximately two-fold between 62.5 μM and 125 μM sodium arsenite but exhibited only a modest, non-significant increase at 250 μM (**Figure 1D**), indicating that SG assembly kinetics approach saturation at approximately 125 μM sodium arsenite.

By contrast, the maximal number of SGs (f) continued to increase with sodium arsenite concentration, rising by approximately 50% between 62.5 μM and 125 μM and by a further ∼10% between 125 μM and 250 μM (**Figure 1E**). This increase occurred despite similar SG assembly rates at the higher sodium arsenite concentrations, indicating that maximal SG number and assembly rate are kinetic parameters that can be modulated independently. Instead, the continued increase in SG number could be explained by progressive shortening of the SG nucleation delay. Cells treated with 250 μM sodium arsenite initiated SG formation approximately 40% earlier than those treated with 125 μM, while cells treated with 125 μM initiated SG formation approximately 40% earlier than those treated with 62.5 μM (**Figure 1F**).

Collectively, these analyses establish a quantitative framework capable of resolving distinct kinetic phases of SG assembly in single cells, providing a sensitive platform for dissecting the mechanisms that regulate SG assembly.

### The eIF4B RRM regulates multiple kinetic phases of SG assembly

Recent studies have shown that the intrinsically disordered regions of eIF4B promote dynamic self-association, suggesting that eIF4B possesses an intrinsic capacity for higher-order assembly.^29^ In addition, our recent work demonstrates that the conserved RNA recognition motif (RRM) regulates RNA-dependent nanometer-scale clustering of the translation initiation machinery^30^, indicating that both the disordered and ordered regions of eIF4B contribute to its higher-order organization. We therefore asked whether disruption of the RRM alters stress granule assembly during cellular stress.

Human eIF4B contains four major functional domains, including a canonical RNA recognition motif (RRM), a DRYG domain that interacts with eIF3, a noncanonical RNA-binding domain, and an eIF4A-binding domain (**Figure 2A**). Because the RRM has recently emerged as a regulator of RNA-dependent higher-order assembly, we examined whether mutations within this conserved domain alter SG formation. To this end, we generated tetracycline-inducible GFP-eIF4B cell lines expressing three separate RRM mutants: F139A (M1), F99A/N102A/R135A (M2), and R135A/K137A/F139A (M3).

**Figure 2.**
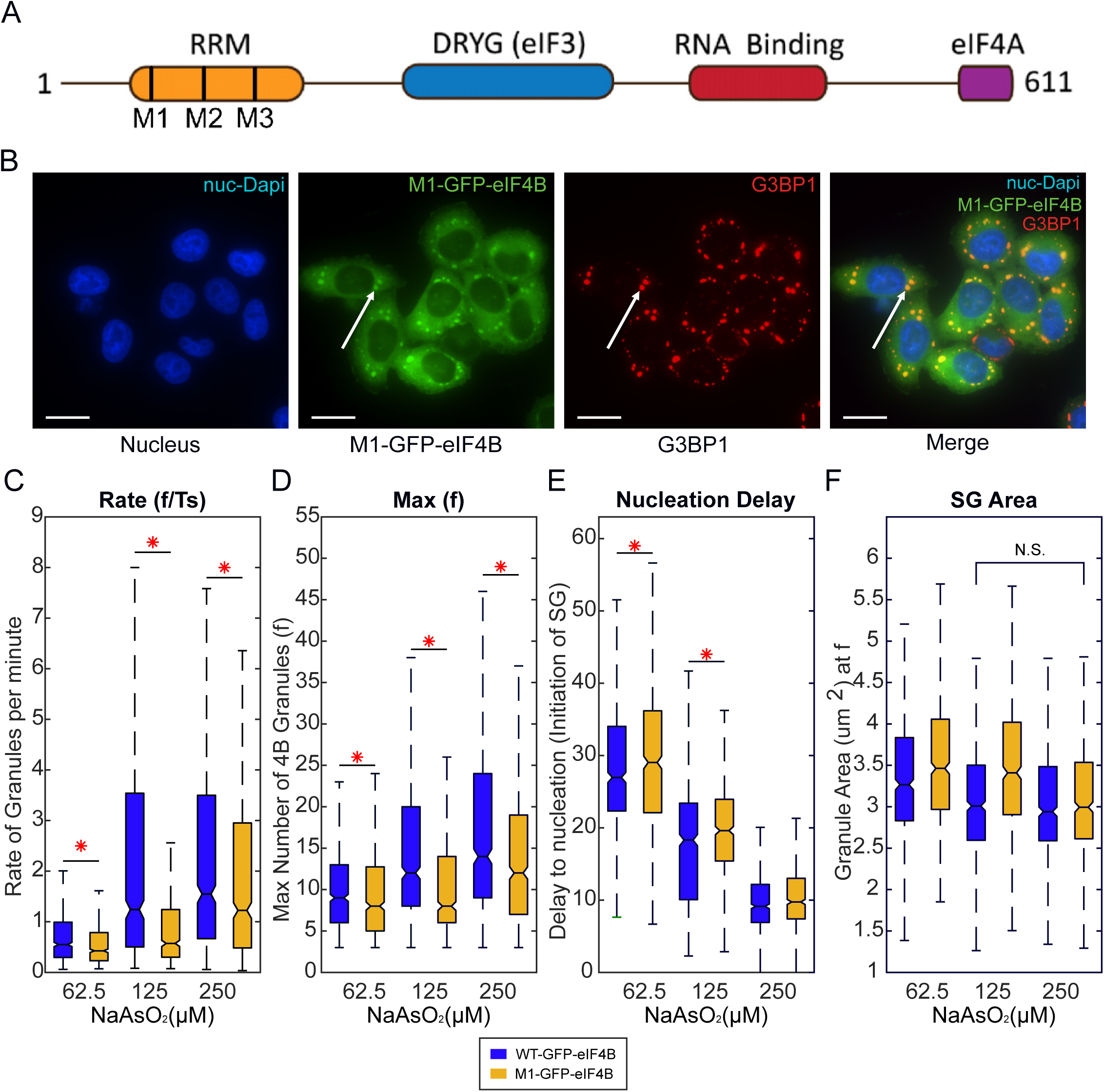
Disruption of the eIF4B RNA recognition motif impairs stress granule assembly. (A) Domain map of the human eIF4B protein includes: RRM domain (amino acids 96-173); DRYG domain (214-327); an arginine-rich motif (ARM) RNA binding domain (367-423); a putative eIF4A binding region (423-611). Mutants were created in the RRM domain: M1 (F139A), M2 (F99A/N102A/R135A), and M3 (R135A/K137A/F139A). (B) Fixed cell images of HeLa cells expressing M1-GFP-eIF4B treated with 125 μM sodium arsenite for two hours. Images from left to right are stained with DAPI to identify the nucleus, GFP-eIF4B, the SG nucleator, G3BP1, and a merged image from all three images to verify GFP-eIF4B in SGs. Scale bars are 25 μm. (C-F) Box and whisker plots showing single-cell WT-GFP-eIF4B and M1-GFP-eIF4B SG kinetics, including **(C)** of rate of formation, **(D)** max number of SGs, **(E)** nucleation delay of SGs, and **(F)** SG area. The following treatment with 62.5 μM, 125 μM, or 250 μM sodium arsenite. Asterisk indicates P<0.005 between groups. Significance was determined via One-Way ANOVA comparison to compare the difference between each treatment group with a Dunnett multi-comparison protocol to account for false discovery rate. n =269 WT and n = 349 M1. Cells per condition pooled from 3 experimental replicates with at least 2 technical replicates in each experiment. See also Figure S2.

Following treatment with 125 μM sodium arsenite for 1 h, formation of GFP-eIF4B puncta was impaired in all mutants compared to WT-GFP-eIF4B (**Figure S2A**). We selected the M1 (F139A) mutant for further analysis because it represents a single amino acid substitution within the conserved RRM that perturbs RNA-dependent higher-order assembly while minimizing structural disruption of the protein. Immunofluorescence for G3BP1 revealed a significant decrease in SGs in M1-expressing cells compared to WT (13.6% reduction in G3BP1 puncta; P = 0.0016) (**Figure S2B**). Because G3BP1 is an essential SG component, this reduction indicates that M1-GFP-eIF4B overexpression impairs SG formation rather than simply preventing mutant eIF4B incorporation into SGs. Consistent with this, M1-GFP-eIF4B colocalized with G3BP1 to a similar extent as WT (Mander’s overlap 0.82 vs. 0.81) (**Figure 2B**).

Western blot analysis showed that after 24 h of tetracycline induction, M1-GFP-eIF4B was expressed at levels comparable to endogenous eIF4B and similar to WT-GFP-eIF4B (**Figure S2C, S1D**). Thus, disruption of SG assembly by the M1 mutant cannot be explained by altered protein expression but instead reflects changes in RRM-dependent function.

We next applied our quantitative kinetic framework to compare SG assembly between M1- and WT-GFP-eIF4B-expressing cells. Disruption of the RRM selectively altered multiple phases of SG assembly. M1-GFP-eIF4B consistently reduced the SG assembly rate (f/Ts) by 15–60% across sodium arsenite concentrations (**Figure 2C**), while the maximal number of SGs formed (f) was reduced by 8–30% (**Figure 2D**). In addition, the SG nucleation delay (Td − Ts/2) was increased at 62.5 μM and 125 μM sodium arsenite but not at 250 μM, suggesting that high-dose sodium arsenite largely saturates SG nucleation kinetics in both WT- and M1-expressing cells (**Figure 2E**).

Finally, we examined whether disruption of the RRM altered SG size in addition to SG assembly kinetics. Across all sodium arsenite concentrations and cell lines, the average SG area ranged from 3.1–3.5 μm², a difference within the imaging resolution (∼1 pixel), indicating no detectable change in SG size (**Figure 2F**). Furthermore, when comparing conditions with similar intrinsic SG assembly rates (WT at 125 μM versus M1 at 250 μM), SG area remained indistinguishable.

Together, these data demonstrate that the conserved RRM of eIF4B is required for efficient SG assembly kinetics. Disruption of the RRM delayed SG nucleation and reduced the rate and extent of SG accumulation, while SGs that did form reached a similar size and retained mutant eIF4B. Thus, the RRM primarily affects early kinetic steps in SG assembly rather than granule growth or incorporation of eIF4B into assembled SGs.

### Quantitative analysis reveals that SG assembly kinetics predict translational repression during oxidative stress

Because disruption of the eIF4B RRM selectively impaired SG assembly kinetics, we next asked whether these defects were associated with altered translational repression during oxidative stress. To address this question, we combined our live-cell imaging assay with a single-cell O-propargyl-puromycin (OPP) incorporation assay that measures nascent protein synthesis. By quantifying OPP incorporation in the same cells that had been tracked throughout SG assembly, we directly related the translational state of individual cells to their complete SG assembly history.

Under basal conditions, M1-GFP-eIF4B cells exhibited approximately 50% lower protein synthesis than WT-GFP-eIF4B cells (**Figure 3A**), indicating that disruption of the RRM impairs translation even in the absence of stress. Surprisingly, despite this lower basal translational activity, oxidative stress produced substantially weaker translational repression in M1-GFP-eIF4B cells than in WT-GFP-eIF4B cells. Following treatment with 125 μM sodium arsenite, protein synthesis was reduced by approximately 74% in M1-GFP-eIF4B cells compared with approximately 93% in WT-GFP-eIF4B cells. This relationship was maintained across all sodium arsenite concentrations examined, with M1-expressing cells consistently exhibiting fewer stress granules together with weaker translational repression than WT cells (**Figure 3B**).

**Figure 3.**
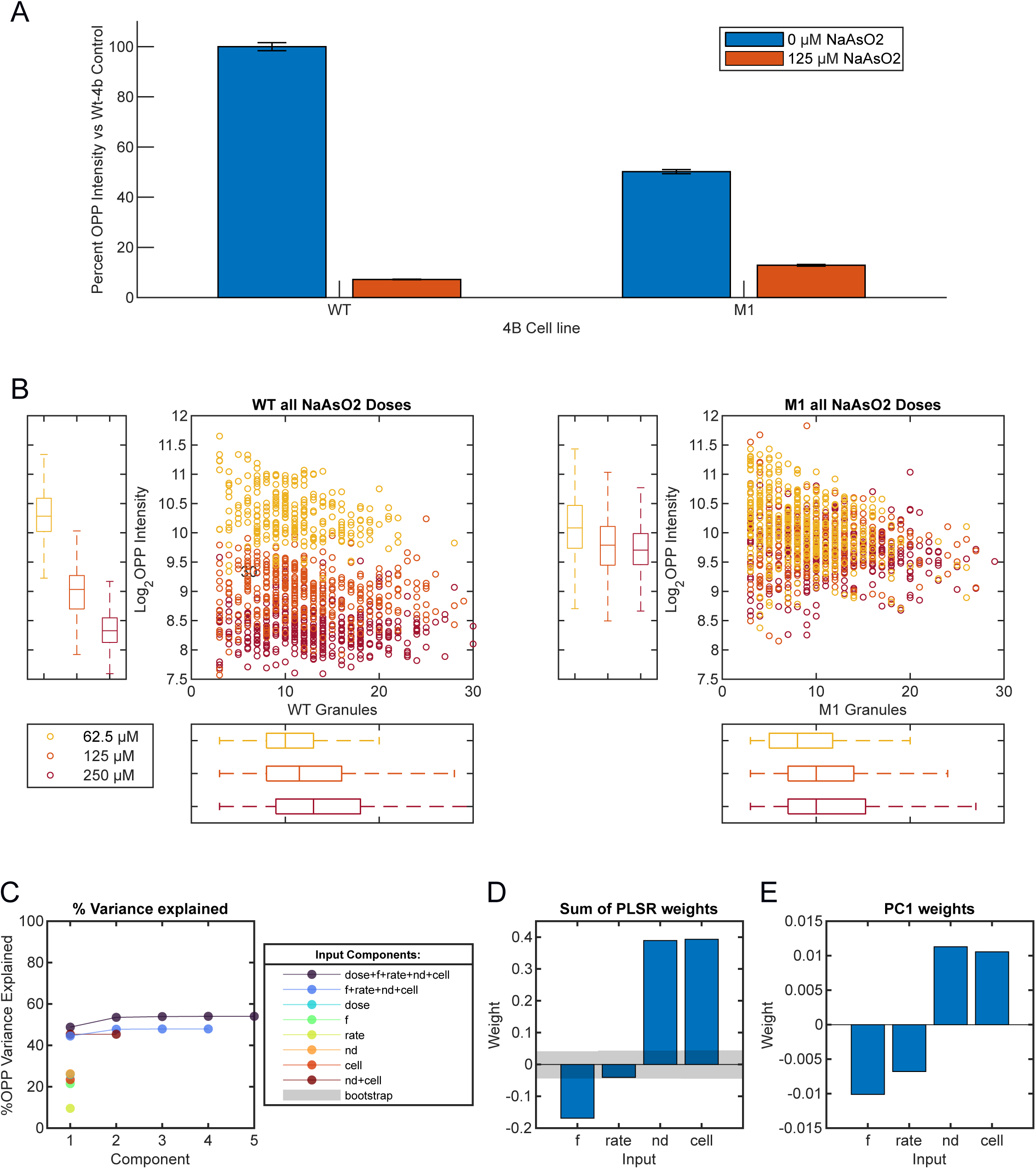
Disruption of the eIF4B RRM impairs stress-induced translational repression. (A) Bar graph with standard error of the mean (SEM) showing the relative % O-propargyl-puromycin (OPP) signal intensity of WT-eIF4B and M1-eIF4B expressing cells treated with vehicle control (blue) or 125 μM Sodium Arsenite (orange) for 2 hours. The p value between the two treatment groups (orange) P<0.005. (B) Box-and-whisker plots showing single-cell Log_2_ O-propargyl-puromycin (OPP) intensity as a function of stress granule (SG) number at the indicated sodium arsenite concentrations. Colors denote arsenite concentration: 62.5 μM (yellow), 125 μM (orange), and 250 μM (red). Left, WT-GFP-eIF4B-expressing HeLa cells; right, M1-GFP-eIF4B-expressing HeLa cells. Each point represents a single cell. (C) Partial least squares regression (PLSR) analysis showing the percentage of variance in Log_2_ OPP explained by models containing different combinations of input variables. Variables included sodium arsenite concentration (dose), maximum SG number (f), SG assembly rate (rate), SG nucleation delay (nd), and cell line identity (cell), as indicated in the legend. For multivariable models, curves show the cumulative variance explained as successive principal components (PCs) are added, whereas single-variable models are represented by a single point. Models incorporating all variables explained 54% of the variance in Log_2_ OPP (purple), while models containing SG kinetic parameters and cell line identity, but excluding arsenite concentration, explained 48% (blue), indicating that SG assembly dynamics account for much of the predictive power of the full model. (D) Relative contribution of individual input variables to prediction of Log_2_ OPP levels. Bars represent the cumulative weights of each input variable across the first four principal components of the PLSR model containing maximum SG number (f), SG assembly rate (rate), SG nucleation delay (nd), and cell line identity (cell) (blue model in Panel C). Gray shaded regions indicate the range expected from 5,000 bootstrap iterations using randomized input data; variables falling outside this range make a significant contribution to the predictive model. Positive and negative weights indicate positive or negative correlations with Log2 OPP, respectively. (E) Weights of input parameters in the first principal component (PC1) of the (f+rate+nd+cell) PLSR model. See also Figure S3.

The combination of real-time SG imaging with single-cell OPP measurements allowed us to directly relate the translational state of individual cells to their complete SG assembly history. We therefore asked whether distinct kinetic phases of SG assembly predict protein synthesis following oxidative stress. Strikingly, SG nucleation delay emerged as the strongest predictor of protein synthesis. Cells exhibiting longer nucleation delays maintained higher levels of protein synthesis, whereas cells that initiated SG assembly rapidly displayed more robust translational repression.

To quantify this relationship, we applied partial least squares regression (PLSR) modeling using SG kinetic parameters (f, rate, nucleation delay), cell line identity (WT or M1), and sodium arsenite concentration as input variables, with single-cell protein synthesis (log₂ OPP intensity) as the output. Together, these variables explained 54% of the variance in protein synthesis (**Figure 3C**). Excluding sodium arsenite concentration reduced predictive power only slightly (to 48%), indicating that SG kinetics and cell line information alone capture much of the observed cell -to-cell variation.

Analysis of predictor contributions revealed that SG nucleation delay, sodium arsenite concentration, cell line identity, and maximal SG number each made substantial contributions to the model, whereas SG assembly rate contributed comparatively little (**Figure 3D–E**). These findings suggest that the timing of SG nucleation, rather than the subsequent rate of SG assembly, is more closely associated with translational repression.

Together, these findings demonstrate that quantitative analysis of SG assembly kinetics provides a powerful framework for understanding translational remodeling during cellular stress. Although several variables contributed to predicting translational output, the timing of SG nucleation was more informative than the subsequent rate of SG assembly, emphasizing the importance of early stages of SG assembly in the cellular response to oxidative stress.

### The eIF4B RRM mutant exhibits selective biochemical defects

The evolutionary conservation of the eIF4B RNA recognition motif (RRM) has long suggested an important role in translation initiation. However, despite more than three decades of biochemical investigation, its precise function has remained unresolved. Early studies demonstrated that disruption of conserved RRM residues reduced stimulation of eIF4A helicase activity while having surprisingly little effect on overall RNA binding, suggesting that the RRM contributes to eIF4B function through a mechanism distinct from bulk RNA affinity.^32^ More recent studies further showed that the RRM is largely dispensable for yeast translation and growth^28^, leaving its physiological role unclear. We therefore asked whether disruption of the human eIF4B RRM selectively alters biochemical activities that could explain its requirement for efficient stress granule assembly.

To quantitatively evaluate the effect of the M1 mutation on helicase stimulation, we measured eIF4A-mediated duplex unwinding in real time using purified eIF4A, eIF4B, and an eIF4G fragment encompassing residues 682–1599. Consistent with both the earlier biochemical studies and our recent single-molecule analysis, the apparent affinity of M1-eIF4B for stimulation of eIF4A-mediated duplex unwinding was reduced approximately two-fold compared with WT-eIF4B (**Figure 4A**). Together, these findings demonstrate that disruption of the RRM reduces, but does not abolish, the ability of eIF4B to stimulate eIF4A helicase activity, indicating that the mutant retains substantial biochemical activity.

**Figure 4.**
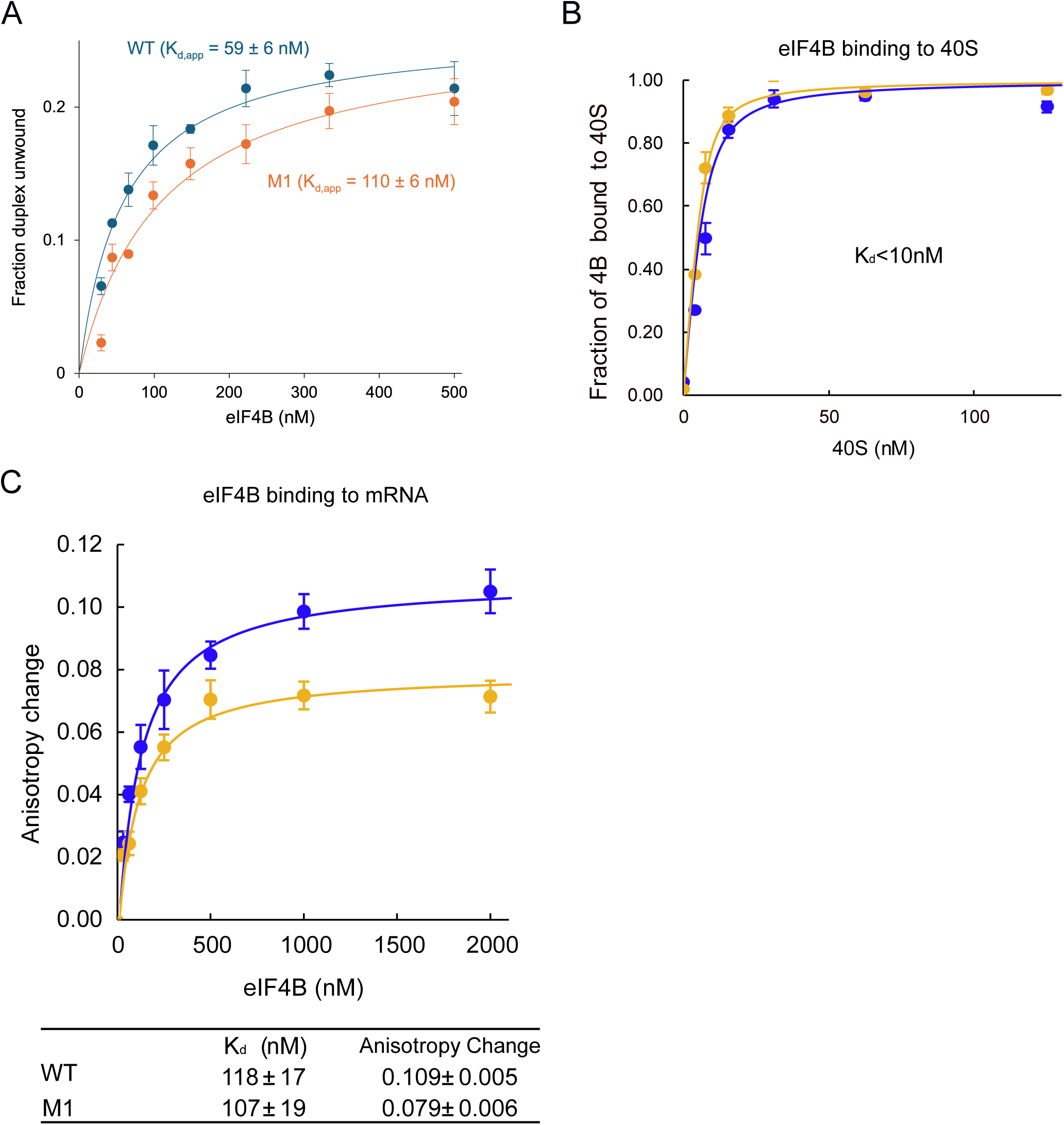
The eIF4B RRM selectively alters biochemical activities. (A) Amounts of ATP-dependent RNA duplex unwound under various amounts WT- or M1-eIF4B in the presence of 0.5 μM eIF4A and eIF4G (682–1599), after 5 min incubation. Apparent dissociation constants (mean ± SEM, n=3) are indicated. (B) Equilibrium binding of fluorescently labeled WT (blue line) or M1 (orange line) eIF4B proteins to the 40S subunit, as measured by an anisotropy assay. The fraction of eIF4B bound at different concentrations of the 40S subunit is shown. The data are the average of at least three trials and error bars indicate the SEM. Equilibrium binding data were fit to determine the equilibrium dissociation constant (K_d_) of eIF4B binding to the 40S subunit, as detailed in the Materials and Methods. (C) Equilibrium binding of fluorescently labeled 42 nt RNA to WT (blue line) or M1 (orange line) eIF4B proteins, as measured by an anisotropy assay. The fraction of eIF4B bound at different concentrations of the 42 nt RNA is shown. The data are the average of at least three trials and error bars indicating the SEM. Equilibrium binding data were fit to determine the equilibrium dissociation constant (Kd) of eIF4B binding to the 42 nt RNA, as detailed in the Materials and Methods. See also Figure S4.

We next asked whether disruption of the RRM alters other core biochemical activities of eIF4B. Previous work has demonstrated that eIF4B binds directly to the 40S ribosomal subunit and that this interaction is largely independent of the RRM^28,33^. We therefore asked whether human eIF4B likewise binds directly to the human 40S ribosomal subunit and whether this interaction is similarly maintained following disruption of the RRM. Using purified human components, WT-eIF4B bound directly to the human 40S ribosomal subunit with high affinity. M1-eIF4B likewise formed a high-affinity complex with the human 40S subunit, although the tightness of the interaction precluded reliable determination of small differences in affinity under these assay conditions (**Figure 4B**). These data indicate that disruption of the RRM does not abolish stable association of human eIF4B with the 40S ribosomal subunit.

Finally, we examined whether disruption of the RRM altered the interaction of eIF4B with RNA. Consistent with previous studies, WT- and M1-eIF4B bound RNA with similar apparent affinities (**Figure 4C**). However, the mutant produced a markedly different maximal fluorescence anisotropy despite exhibiting little change in overall RNA affinity. These findings indicate that disruption of the RRM alters the way eIF4B engages RNA rather than simply reducing its affinity for RNA.

Collectively, these findings indicate that disruption of the RRM selectively alters the RNA-dependent activities of eIF4B while preserving stable interactions with both the 40S ribosomal subunit and RNA. Although the mutant exhibits only a modest reduction in helicase stimulation, it displays altered RNA engagement, consistent with our recent demonstration that the same mutation impairs RNA-dependent nanometer-scale clustering of the translation initiation machinery at concentrations well below the threshold for phase separation.^30^ Together, these findings support a model in which the RRM promotes productive RNA-dependent organization of eIF4B rather than serving as a primary determinant of ribosome or RNA binding.

## Discussion

In this study, we established a quantitative single-cell framework for resolving distinct phases of stress granule (SG) assembly during oxidative stress. By combining live-cell imaging with kinetic modeling, we quantified discrete kinetic phases of SG assembly, including nucleation delay, assembly rate, and maximum SG number. This approach extends beyond conventional endpoint analyses by monitoring the delay, rate, and extent of SG assembly independently, providing a general framework for investigating the dynamic regulation of stress granule assembly in living cells. Furthermore, coupling this kinetic analysis with single-cell measurements of protein synthesis allowed us to relate the temporal history of SG assembly to translational state of each cell. This integrated approach therefore provides a means to investigate how SG dynamics are associated with translational remodeling.

Our findings also provide new insight into the long-standing question of the biological function of the conserved eIF4B RNA recognition motif (RRM). Although eIF4B has long been recognized as an essential cofactor for eIF4A, the function of its RRM has remained surprisingly elusive. Early biochemical studies demonstrated that disruption of conserved RRM residues reduced stimulation of eIF4A helicase activity despite having relatively little effect on overall RNA binding, while subsequent genetic studies in yeast showed that the RRM is largely dispensable for translation initiation and cell growth.^28,32^ Together, these observations suggested that the RRM serves a regulatory rather than an essential role in eIF4B function, although the nature of this regulatory activity remained unclear. The evolutionary conservation of the RRM from yeast to humans, despite its apparently modest contribution to basal translation, suggests that its principal function is most apparent during periods of rapid reorganization of the translation initiation machinery.

Recent studies have shown that the intrinsically disordered regions of eIF4B promote dynamic self-association^29^, while the conserved RRM promotes RNA-dependent nanometer-scale clustering of the translation initiation machinery (RPCs).^30^ Viewed in the context of the present study, these findings suggest that the conserved role of the RRM is not simply to increase RNA binding affinity or ribosome association, but rather to promote productive higher-order organization of the translation initiation machinery. During cellular stress, this organizational function may facilitate efficient SG assembly and the associated translational remodeling required for cellular adaptation.

Our biochemical analyses are consistent with this emerging model. Disruption of the RRM reduced, but did not abolish, stimulation of eIF4A helicase activity while preserving high-affinity association with both RNA and the 40S ribosomal subunit. Perhaps the most striking biochemical consequence of the mutation was instead an alteration in the way eIF4B engaged RNA despite little change in overall RNA affinity. In our recent work, the same mutation impaired nanometer-scale RPC formation by the translation initiation machinery both *in vitro* and in living cells, producing smaller and fewer RPCs than WT eIF4B.^30^ Thus, rather than simply mediating individual interactions with RNA or the ribosome, the RRM may promote productive RNA-dependent interactions that facilitate higher-order organization of initiation complexes.

It is tempting to speculate that the higher-order organization of the translation initiation machinery promoted by eIF4B may also contribute to SG nucleation. The finding that the same RRM mutation impairs RPC formation while delaying SG nucleation and slowing SG assembly raises the possibility that RPCs contribute to the early stages of SG formation, perhaps by providing a pool of intermediate RNP assemblies from which SGs can nucleate. Such a model would be consistent with emerging evidence that RNP condensates can arise through multivalent interactions among smaller RNA–protein assemblies, and that formation of higher-order oligomers can precede macroscopic condensation.^29^ Whether RPCs represent direct intermediates in the SG assembly pathway, however, remains to be determined. The selective nature of these biochemical defects may explain why disruption of the RRM has a relatively modest effect on basal translation yet produces pronounced defects in SG assembly and translational remodeling during cellular stress, when efficient coordination of multiple translation initiation components is likely to become particularly important.

Our quantitative analysis also demonstrates that SG assembly is not a single event, but rather comprises multiple kinetic phases that can be independently regulated. Disruption of the eIF4B RRM delayed SG nucleation, slowed SG assembly, and reduced the total number of SGs formed, while having little effect on the final size of mature SGs. The selective effects on nucleation and assembly, but not mature SG size, suggest that distinct molecular mechanisms contribute to different stages of SG formation rather than SG growth or maturation. More broadly, our findings illustrate how quantitative kinetic analyses can reveal mechanistic insights that are not apparent from conventional endpoint measurements alone.

Our findings also provide new insight into the relationship between SG assembly and translational control. Although SGs are widely regarded as a consequence of translational repression, the temporal relationship between SG assembly and changes in protein synthesis has remained difficult to investigate. Rather than relying on static endpoint measurements, this integrated approach enabled the translational state of individual cells to be interpreted in the context of their prior SG assembly history, revealing that defects in SG assembly are closely associated with altered translational remodeling during oxidative stress. This relationship provides a quantitative basis for investigating how dynamic changes in SG assembly relate to translational adaptation under diverse physiological conditions.

Collectively, our findings support a model in which the conserved RNA recognition motif of eIF4B promotes productive higher-order organization of the translation initiation machinery rather than simply strengthening individual interactions with RNA or the ribosome **(Figure 5)**. This organizational activity may become particularly important during cellular stress, when translation initiation complexes must be rapidly reorganized to support SG assembly and translational adaptation.

**Figure 5.**
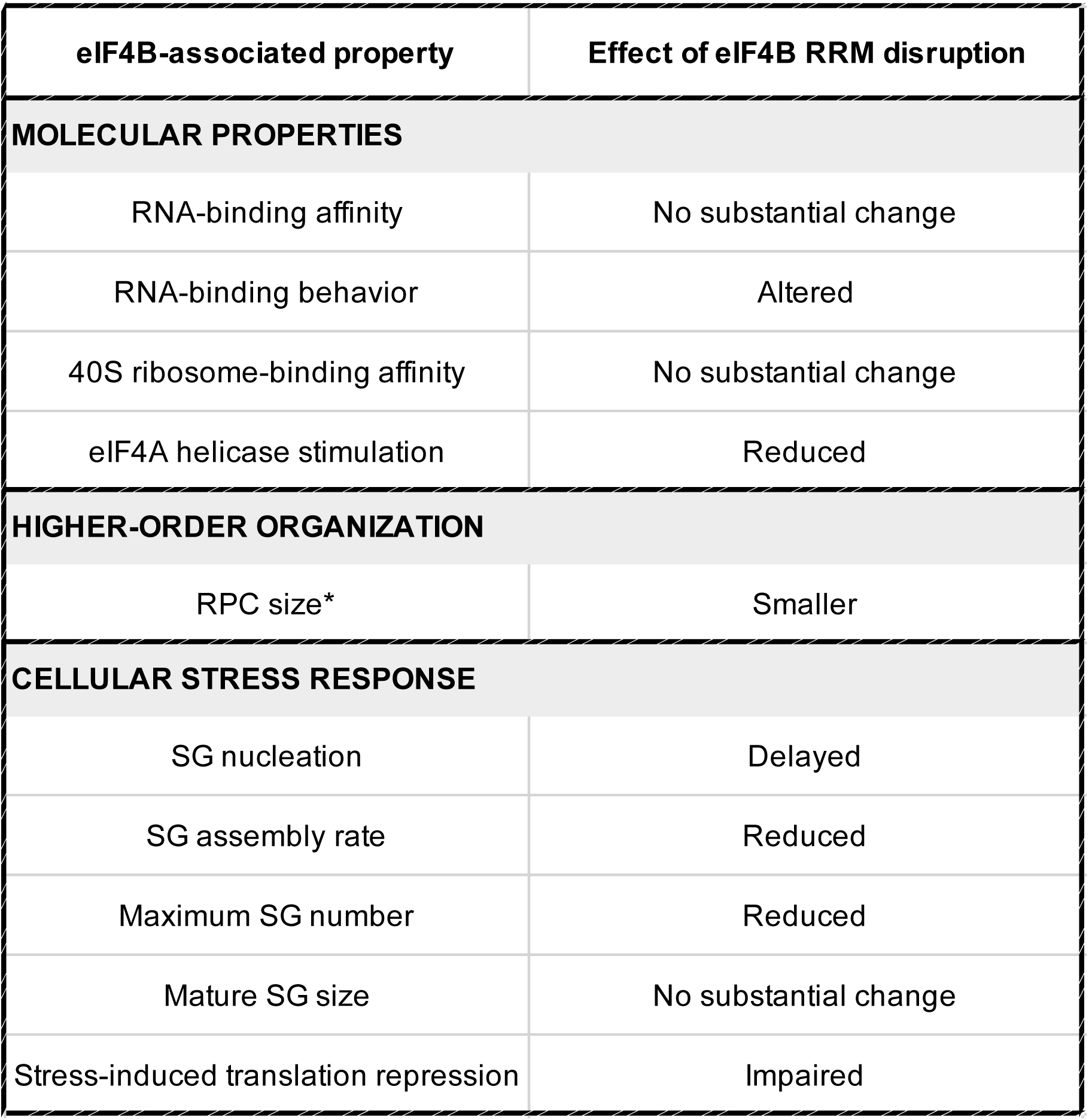
Summary of the effects of eIF4B RRM disruption. The table summarizes the effects of disrupting the eIF4B RNA recognition motif (RRM) on molecular properties of eIF4B, higher-order organization of the translation initiation machinery, and the cellular response to oxidative stress. RRM disruption has little effect on overall RNA-binding affinity or 40S ribosome-binding affinity, but alters RNA-binding behavior and reduces stimulation of eIF4A helicase activity. *The same RRM mutation produces smaller RNA-Protein-Clusters (RPCs), as described in Ref. 30. During oxidative stress, RRM disruption delays SG nucleation, reduces the rate of SG assembly and maximum SG number, and impairs stress-induced translational repression, while having little effect on mature SG size. These observations support a model in which the eIF4B RRM promotes higher-order organization of the translation initiation machinery and suggest that this function may contribute to efficient SG assembly and translational remodeling during cellular stress.

Similar organizational mechanisms may operate more broadly in cellular contexts requiring rapid translational reprogramming. More generally, our findings suggest that higher-order organization of the translation initiation machinery represents an additional layer of translational regulation, while quantitative kinetic analysis of SG assembly provides a means to investigate these dynamic processes across diverse physiological and pathological settings.

## Acknowledgements

We wish to thank Nancy Villa for her contributions to the early stages of this work, including assistance with cell line development, and for many insightful discussions. We thank Michael Pargett for his discussions on PLSR. We would like to thank Drs. Elena Dobrikova and Matthias Gromeier from Duke University Medical Center for the gift of the HeLa Flp-In T-Rex host cell line.

## Author Contributions

J.B. and C.S.F. conceived and designed the study and led the project. N.L.D. conceived the live-cell imaging and OPP experiments, developed the image analysis and data-processing pipelines, and performed the analysis and model fitting of the live-cell and immunofluorescence data. J.B., N.L.D., E.K., and M.S. performed experiments. K.B. generated the GFP-eIF4B cell line. J.B. and N.L.D. prepared Figures 1–3 and Supplemental Figures 1–4, and M.S. prepared Figure 4. J.G.A. provided guidance on experimental design. J.B. and N.L.D. wrote the manuscript with input and editing from all authors.

## Funding and additional information

This work was supported by the NIH grant R35GM152137 to CSF, and R01HL151983 and R35GM139621 to JGA. NLD was supported by the UC Davis Lung Center T32 - NIH HL007013. The content is solely the responsibility of the authors and does not necessarily represent the official views of the National Institutes of Health.

## Methods

### Key Resources Table

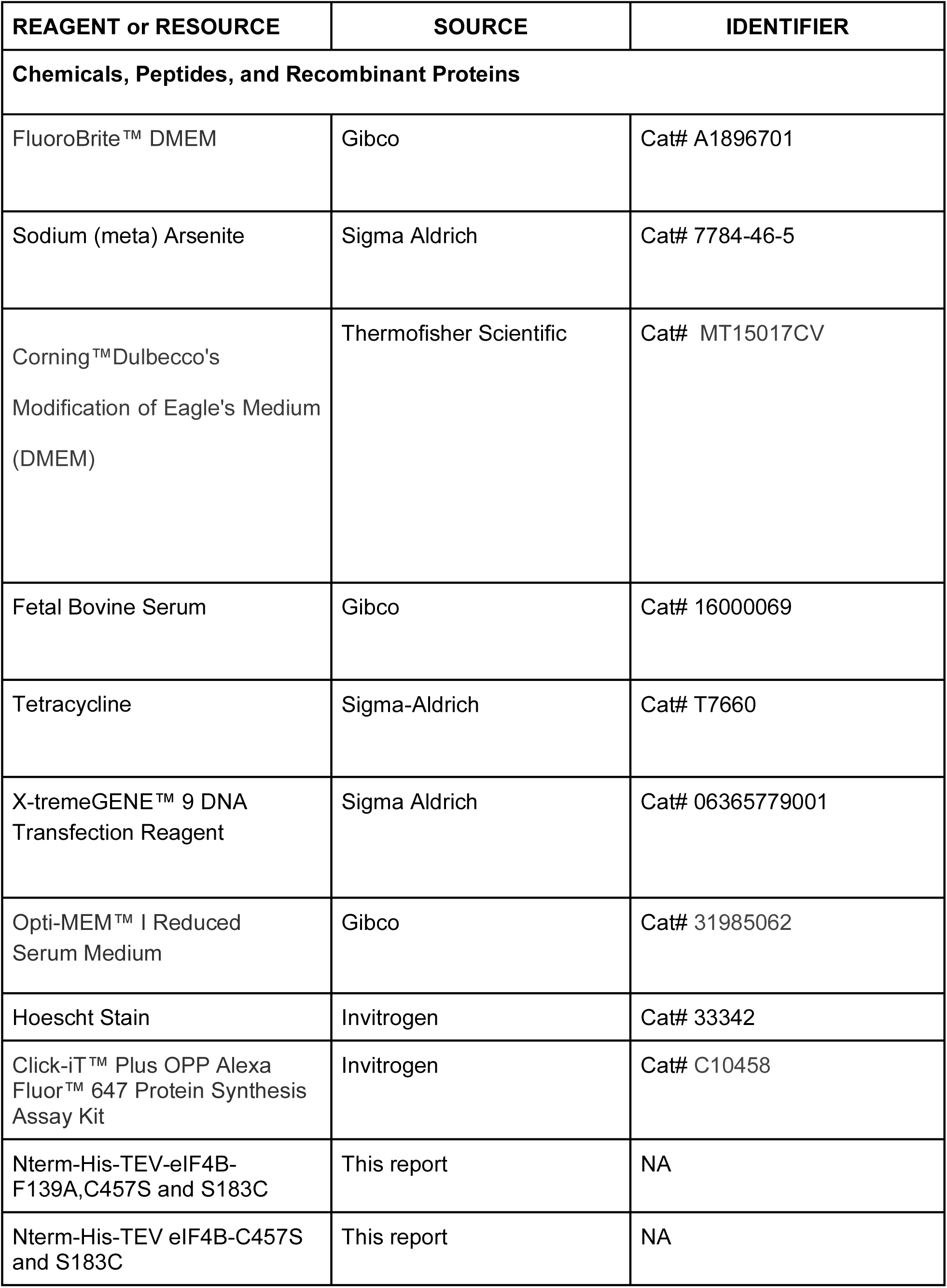
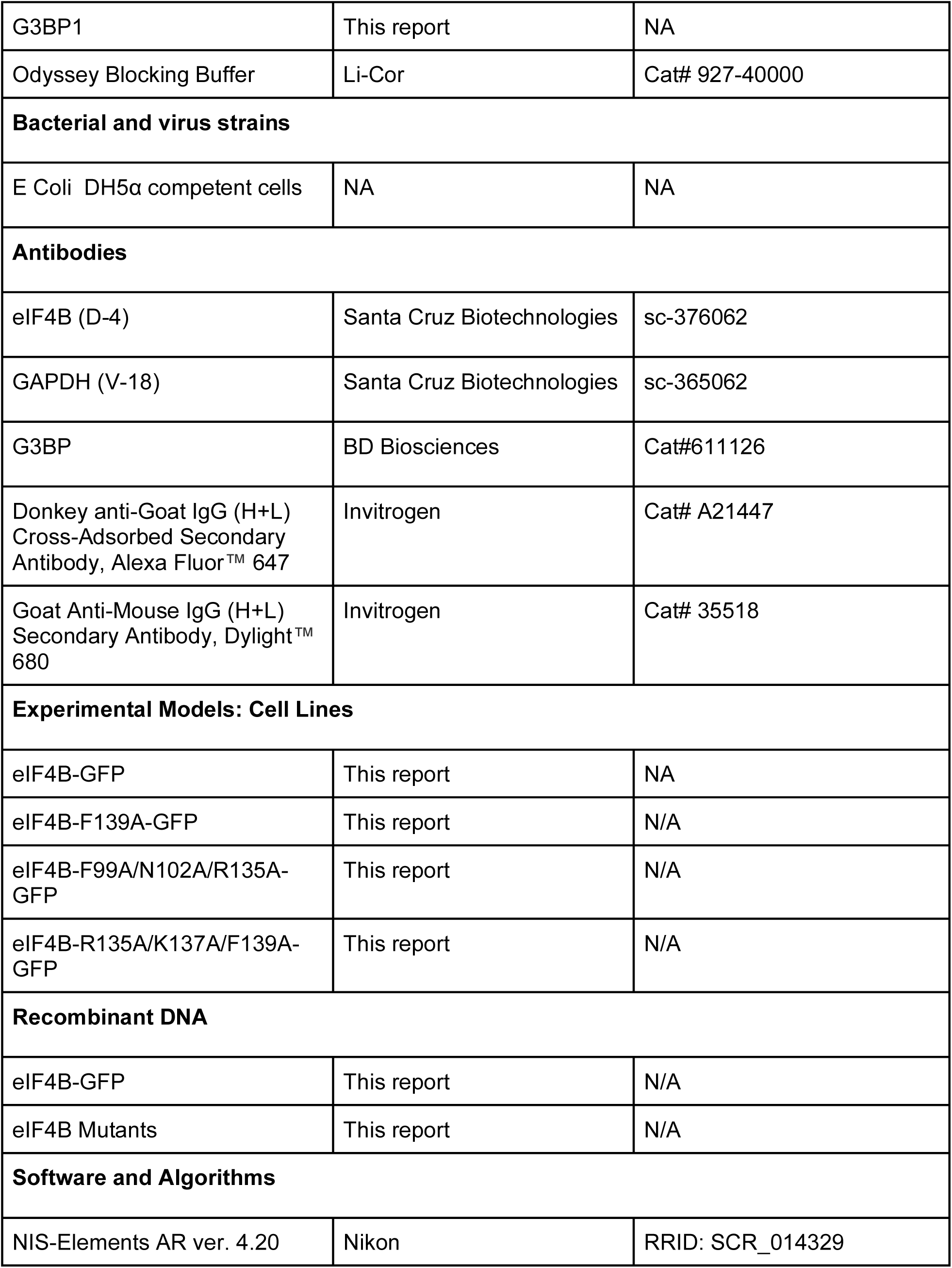
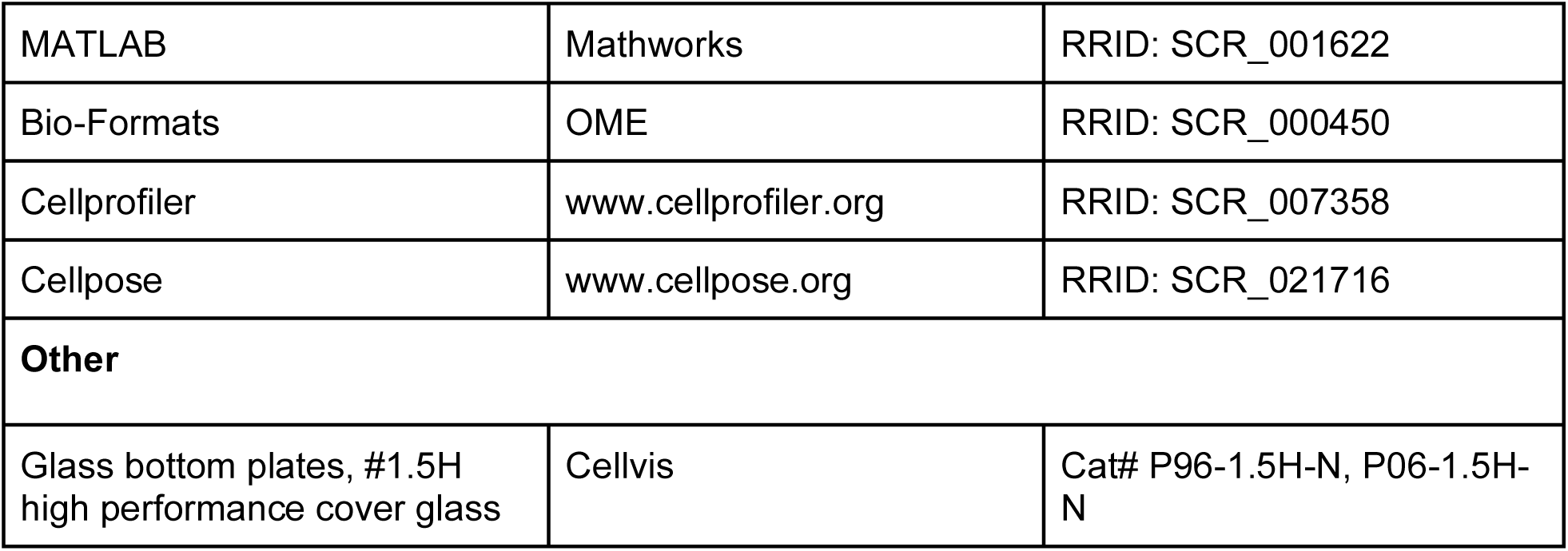

| REAGENT or RESOURCE | SOURCE | IDENTIFIER |
| --- | --- | --- |
| **Chemicals, Peptides, and Recombinant Proteins** |  |  |
| FluoroBrite™ DMEM | Gibco | Cat# A1896701 |
| Sodium (meta) Arsenite | Sigma Aldrich | Cat# 7784-46-5 |
| Corning™Dulbecco's<br>Modification of Eagle's Medium<br>(DMEM) | Thermofisher Scientific | Cat# MT15017CV |
| Fetal Bovine Serum | Gibco | Cat# 16000069 |
| Tetracycline | Sigma-Aldrich | Cat# T7660 |
| X-tremeGENE™ 9 DNA<br>Transfection Reagent | Sigma Aldrich | Cat# 06365779001 |
| Opti-MEM™ I Reduced<br>Serum Medium | Gibco | Cat# 31985062 |
| Hoescht Stain | Invitrogen | Cat# 33342 |
| Click-iT™ Plus OPP Alexa<br>Fluor™ 647 Protein Synthesis<br>Assay Kit | Invitrogen | Cat# C10458 |
| Nterm-His-TEV-eIF4B-F139A,C457S and S183C | This report | NA |
| Nterm-His-TEV eIF4B-C457S<br>and S183C | This report | NA |
| G3BP1 | This report | NA |
| Odyssey Blocking Buffer | Li-Cor | Cat# 927-40000 |
| **Bacterial and virus strains** |  |  |
| E Coli DH5α competent cells | NA | NA |
| **Antibodies** |  |  |
| eIF4B (D-4) | Santa Cruz Biotechnologies | sc-376062 |
| GAPDH (V-18) | Santa Cruz Biotechnologies | sc-365062 |
| G3BP | BD Biosciences | Cat#611126 |
| Donkey anti-Goat IgG (H+L)<br>Cross-Adsorbed Secondary<br>Antibody, Alexa Fluor™ 647 | Invitrogen | Cat# A21447 |
| Goat Anti-Mouse IgG (H+L)<br>Secondary Antibody, Dylight™<br>680 | Invitrogen | Cat# 35518 |
| **Experimental Models: Cell Lines** |  |  |
| eIF4B-GFP | This report | NA |
| eIF4B-F139A-GFP | This report | N/A |
| eIF4B-F99A/N102A/R135A-GFP | This report | N/A |
| eIF4B-R135A/K137A/F139A-GFP | This report | N/A |
| **Recombinant DNA** |  |  |
| eIF4B-GFP | This report | N/A |
| eIF4B Mutants | This report | N/A |
| **Software and Algorithms** |  |  |
| NIS-Elements AR ver. 4.20 | Nikon | RRID: SCR_014329 |
| MATLAB | Mathworks | RRID: SCR_001622 |
| Bio-Formats | OME | RRID: SCR_000450 |
| Cellprofiler | www.cellprofiler.org | RRID: SCR_007358 |
| Cellpose | www.cellpose.org | RRID: SCR_021716 |
| **Other** |  |  |
| Glass bottom plates, #1.5H<br>high performance cover glass | Cellvis | Cat# P96-1.5H-N, P06-1.5H-N |

## Resource availability / Lead contact

For further information and any requests for resources and reagents, please contact Drs. Christopher Fraser or John Albeck.

## Materials availability

Plasmids and cell lines generated in this study are available from lead contacts.

### Data and Code availability

Raw NIS elements image data can be made available upon request, but were not uploaded to the submission portal due to their large size.

All code used to import cellprofiler data into Matlab, filter and process live-cell or IF data, pool experimental replicates, align live-cell to IF data, fit data models, produce figures, and perform statistical analysis are available on github at: https://github.com/Albeck-Lab/SG_4B

Figures 1-3 have html output files which can be read on any device, showing all code used for each figure at: https://github.com/Albeck-Lab/SG_4B/tree/main/Paper_Figures/html

### Plasmid Construction

To create stable inducible cell lines, wild-type human eIF4B with a N-terminal EGFP tag was subcloned into the pcDNA5/FRT/TO vector (Invitrogen) using BamHI and XhoI restriction sites. Point mutations in the eIF4B RRM were synthesized by Genscript and subcloned into the above plasmid using ClaI and BstBI restriction sites, as described in the following vectors: M1 (F139A), M2 (F99A, N102A, R135A), M3 (R135A, K137A, F139A). For *in vitro* experiments, eIF4B with an N terminal His-TEV tag was expressed from a derivative of the pET28c vector, as described previously.^34^ To fluorescently label eIF4B, the single native cysteine residue was mutated to serine (C457S) and a new cysteine was introduced (S183C) using site-directed mutagenesis with the following primers:

C457S F primer GAAGATTCCCACTCACCGACGTCGAAACCG,

C457S R primer TGAGTGGGAATCTTCTTCTTTATTCAGGGT,

S183C F primer ACGATCGTTGCTTTGGTCGCGATCGTAACCG

S183C R primer CAAAGCAACGATCGTCGCGGTCTTTATCTTGG

To test the M1 mutant in fluorescence assays this mutation was also added using site-directed mutagenesis with the following primers:

4B RRM F139A F primer TGAAAGGTGCAGGCTATGCAGAATTCGAAG,

4B RRM F139A R primer AGCCTGCACCTTTCAGACGTTCCGGATTAG.

All primers were purchased from Integrated DNA Technologies. All plasmid sequences were confirmed by Sanger sequencing (Azenta Life Sciences and Quintara Biosciences).

### Cell Culture and Imaging Medium

The HeLa Flp-in TRex cell line was a generous gift from Drs. Elena Dobrikova and Matthias Gromeier (Duke University Medical Center). All HeLa cell lines were maintained in DMEM supplemented with 10% fetal bovine serum (FBS) and grown at 37°C in 5% CO_2_. For all imaging experiments, media was replaced with Gibco FluoroBrite DMEM supplemented with 2% FBS for at least 1 hour prior to imaging (or first treatment) to minimize background fluorescence (referred to as ‘imaging medium’).

### Creating stable cell lines

All transfections consisted of the gene of interest inserted into a pcDNA5/FRT/TO plasmid and the plasmid pOG44. All transfections are carried out in Opti-MEM Reduced Serum Media (Thermo Fisher) supplemented with 3% FBS according to manufacturer’s guidelines with minor changes. For each transfection, 1.5–3.0 μg of pcDNA5/FRT/TO plasmid and 1 μg of POG44 is diluted into 100 µL of Opti-MEM media. For each transfection, 6 µL of X-tremeGENE9 DNA Transfection Reagent (Roche) was diluted in 100 µL of Opti-MEM media and incubated for 5 minutes at room temperature. The two incubations were then combined to create a DNA-lipid mixture that was incubated for 30 minutes at room temperature. The transfection was then added to the cell line and incubated for 5 hours. Stable cell lines were generated with Blasticidin Hygromycin selection using a 6-well plate according to manufacturer’s guidelines.

### Live-Cell imaging and Immunofluorescence Microscopy

For all live-cell, immunofluorescence, and OPP assays, GFP-eIF4B HeLa cells were seeded at 10,000 - 20,000 cells per well in a 96-well glass bottom imaging plate (CellVis - P96-1.5H-N) 48 hours prior to the experiment. Expression of GFP-eIF4B was induced by the addition of tetracycline at 0.1 μg/μL final concentration for 2–24 hours prior to the experiment start as indicated (24 hours induction for data in Figures 1-3, Sup Figure 1 A&B, Sup Figure 2 A&B; 0-24 hours of induction for data in Sup Figure 1C&D, Sup Figure 2C, and Sup Figure 3). Cells were washed once with 200 µL imaging medium per well, then incubated in 200 µL imaging media per well for 1 hour prior to the start of experiments and maintained in this medium until the conclusion of the experiment. Sodium Arsenite was added to the cells via the addition of a 10 µL of a 21X stock (of the final concentration desired) solution in imaging medium.

Time-lapse wide-field microscopy was performed using a Nikon (Tokyo, Japan) 40X/0.95 NA Plan Apo objective on a Nikon Eclipse Ti-e or Ti2-e inverted microscope, equipped with a Lumencor SPECTRA X or SPECTRA iii light engine, and Andor Zyla 5.5 scMOS or Teledyne Photometrix Kinetix scMOS camera. Fluorescence filters used are: <u>DAPI</u> (<u>Chroma filters:</u> Excitation: ET395/25x, Mirror: T425lpxr Emission: ET460/50), GFP (49002, Chroma), Orange (Ex: ET546/22x, Mirror: T560lpxr, Em: ET572/23m), and Cy5 (49006, Chroma). Cells were imaged at 2–3 minute intervals with relative powers ranging from 5-35%, and exposure times ranging from 100–400 ms depending on expression and fluorescence. During Live-Cell experiments cells were imaged at 2–3 minute intervals and maintained at 37°C with 5% CO_2_ on the microscope via OKO systems.

### Immunofluorescence

After performing the SG assay, the cells were fixed and immunofluorescent staining was performed for G3BP1 via the following protocol. Cells were permeabilized and fixed by the addition of 16% PFA (for a final concentration of 2% PFA) directly to the media and incubated for 10 minutes at room temperature in a fume hood. The PFA/media solution was removed by pipetting, and cells were washed once with 150 µL of 1X PBS per well with gentle rocking for 10 minutes. The PBS was removed and 100 µL MeOH was added per well and incubated for 10 minutes in the fume hood. MeOH was removed via pipetting and the cells were washed three more times by gentle rocking with 100 µL PBS per well for 5 minutes each wash. After the washes, the PBS was removed and 100 µL of Odyssey blocking buffer was added per well and incubated with gentle rocking at room temperature for 60 minutes. Then, the blocking buffer was removed and 100 µL of blocking buffer containing a 1:250 dilution of G3BP1 Primary Antibody was added to each well (G3BP1 Primary Antibody - BD Transduction mouse anti human G3BP1). The plate was wrapped in aluminum foil to reduce light exposure and incubated for 1 hour at room temperature on a rocking plate, then washed three times with 100 µL PBS-T (0.1% Tween-20 in PBS) for 5 minutes at room temperature. The PBS-T was removed via pipetting, then 100 µL of blocking buffer containing a 1:1000 dilution of Secondary Antibody and a 1:10000 dilution of Hoechst (for nuclear staining) was added to each well (Alexa Fluor 647). Still covered, the plate was incubated at room temperature for 1 hour on a rocking plate, then washed three times with 100 µL PBS-T (per well) for 5 minutes on a rocking plate. Finally, the PBS-T was aspirated and 200 µL PBS was added to each well and imaged as described above.

### Single cell protein synthesis estimation by O-proparagly-puromycin (OPP)

To measure global protein synthesis, cells were labeled with the puromycin analog O-propargyl-puromycin (OPP) at 10 μM final concentration for 30 minutes before the end of live-cell imaging (1.5 hours after stress induction) experiments. Wells were fixed with 2% PFA solution, washed with PBS, methanol permeabilized, washed with PBS twice (as described in detail above in l ive cell imaging section), then incubated with click chemistry reaction buffer (10 μM Azide dye, 4 mM CuSO4, 50 mM Ascorbic acid in 100 mM Tris Buffer pH 8.5 and Alexa 647 Azide dye) for 1 hour per the manufacturer’s directions. Finally, cells were washed twice with PBS, then imaged.

### Image Analysis, SG Modeling, and PLSR

To extract live-cell SG data, first the time lapse images are aligned over time using the NIS elements “align images” software package. This is done to eliminate any potential shifts in the image field that could be introduced during experimental treatment spikes which could cause breaks in cell/granule tracking in later processing steps. Once aligned, live-cell and IF data are fed into a custom cellprofiler 4 pipeline.^35^ This pipeline includes cell identification via the built-in cellpose model cyto2 and isolation of these identified cells. Stress granules were then identified in the cells using a custom cellpose model we trained to identify stress granules (called SGI_High_Contrast; available on our github).^36,37^ Where applicable (live-cell data), cells were tracked over time using the “follow neighbors’’ track objects module, and verified proper cell and granule identification using the overlay outlines module. Cell and granule intensity, size, shape, location, and counts were exported to csv files placed into folders per image field (XY) for that experiment. For immunofluorescence (IF) images (i.e. G3BP1) a similar pipeline was used that did not track the objects over time.

Once the Cell Profiler pipeline extracted the SG data, the csv files were iteratively opened in Matlab, where the metadata from that experiment (including which cell line and treatments were in each XY) are appended to the Live-Cell or IF data. Live-cell data was filtered to ensure quality by requiring a minimum amount of time the cells were tracked to be at least 1 hour. Where applicable, IF data was aligned to the Live-Cell data using the coordinates of the cells in the last frame of the movie and a KNN-search for cells within a given radius. This data was saved per experiment in a data object and performed via the function SG_Datahandler.

To extract SG kinetics from Live-Cell images, data was fed into a custom function called convertDatalocToModelFit. This function loops over the experimental datasets provided and for each cell uses nonlinear least squares to fit the following equation to the SG number of that cell using Matlab’s built in ‘fit’ function. **“f-f/(1+exp(((xdata-Td)/(Ts/4))))”.** Figures 1 and 2 used data aggregated from three experimental replicates that had at least two technical replicates each.

Because GFP-eIF4B granules first appear, reach a maximum number, then begin to aggregate over time, this can cause poor model fitting. To circumvent this, each cell’s data was truncated to include the point of treatment until 1 time point (3 minutes) after reaching that cell’s maximum SG count (f). To accomplish this a fixed f was fed into the model along with the SG count, and xdata is the timepoints (in minutes) of the data included. The fit function then used non-linear least squares regression to solve for Td (time from treatment to half max SGs) and Ts (time from first SG appearance to max SGs). Fit models with an R^2^ of less than 0.8 were filtered out of the data.

Partial least squares regression (PLSR) was performed using previously published methodology.^38^ Briefly, a PLSR model which could explain the variance in single-cell Log_2_ OPP between cells expressing either WT- or M1-GFP-eIF4B was solved using the single-cell the metrics of SG kinetics (maximum number of SGs formed, rate of SG formation, SG nucleation delay), sodium arsenite concentration, and the type of GFP-eIF4B expressed. GFP-eIF4B expression was designated for PLSR input by assigning a 0 value to WT-GFP-eIF4B expressing cells and a 1 value to M1-eIF4B expressing cells. All inputs were z-score normalized within the parameter before being fed into the PLSR function. To validate that the PLSR model variance explained was significant, it was compared to a “scrambled” PLSR model. A scrambled PLSR model took the same data and randomly reassigned the cell’s input data, then was solved. To ensure consistency, the scrambled PLSR model was performed 5,000 times. Due to the variability in relative intensity of OPP staining from run to run (e.g. OPP intensities for all cells are 20% higher in one experimental replicate compared to another), data in figure 3 is from one representative experiment (that was also used in figures 1 and 2), with the trends and significance indicated being found in replicate experiments as well. If experimental replicates are pooled by first z-score normalizing the Log_2_ OPP intensities, the OPP data across experimental replicates follow similar trends and maintain statistical significance, suggesting reproducibility between experimental replicates.

### Cell extracts and Western Blotting

Cells were seeded at 1.5×10^6^ in a 10 cm dish. When cells reached 80% confluent, cells were scraped and placed in a lysis buffer. Each cell pellet was lysed in 200 μL of lysis buffer containing 20 mM HEPES, pH 7.5, 10 mM KCl, 1 mM DTT, 1% NP40, 5 mM Mg Acetate and 1x protease inhibitor. Lysis was done using a vortex mixer, in which cells were vortexed for 30 seconds and rested on ice for 1 minute for a total of 10 times. SDS PAGE gels were transferred to Immobilon-FL polyvinylidene fluoride (PVDF) membrane. An overnight transfer was performed at 35V. PVDF membranes were incubated with primary antisera specific to eIF4B (Santa Cruz Biotechnology Lot #B2322) at 1:1000 in Tris-buffered saline with 0.1% Tween 20 (TBST) overnight at 4 °C. PVDF was washed three times for 10 min in TBST, followed by 1 hour incubation at room temperature with goat anti-mouse secondary antibody DyLight 680 (Invitrogen/Thermo Fisher Scientific PI35518). Lastly, Western blots were scanned on an Azure Sapphire Biomolecular Imager (Azure Biosystems).

### Mathematical Model for SG Kinetics

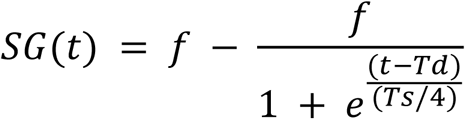

The equation used to model the kinetics of granule formation was modified from the equation in Albeck *et. al.* which was used to describe the cleavage of caspase substrates in individual cells.^31^ Whereby, **SG(*t*)** is the number of stress granules at a given time (in minutes), ***f*** is the maximum number of granules formed in that cell, **Ts** is the time from first SG nucleation event to the time when the max number of granules formed (i.e. when the cell reaches its *f*), and **Td** is the time from treatment (in our case, with sodium arsenite) to when the cell reaches half of the max number of granules it will form (i.e. *f* / 2). From these parameters **f/Ts** which is the rate of granule nucleation (i.e. from first nucleation event until reaching the max number of granules), and **Td- (Ts/2)** which is the time from treatment to the first measured SG in the cell (also referred to as nucleation delay) was calculated.

### Biochemical sample preparations

Human eIF4A1 and eIF4G1 (residues 682-1599) were purified from overexpressing E. coli as described previously.^23,34^ The 40S subunit was purified from HeLa cell extract as described previously.^39^ The 42-nt unstructured RNA with CAA repeat was transcribed and labeled with fluorescein at the 3’-end as described previously.^40^ N-terminal His_6_-tagged WT- and M1-eIF4B were expressed in *E. coli*, with their sequence essentially identical to that previously expressed in sf9 insect cells^23^, but have C457S and S183C mutations for a labeling purpose in addition to F139A in M1-eIF4B. eIF4B was purified through Ni-NTA, Heparin, Mono S, and Superdex 200 prep grade columns, and labeled at C183 with fluorescein-5-maleimide as described previously.^40^ Quality of purified eIF4B proteins and their labeled forms were analyzed in SDS gels (Fig. S4A).

### Helicase Assay

The helicase assay was carried out as described previously.^34,41,42^ The reaction contained 50 nM RNA duplex substrate, 0.5 μM eIF4A1 and eIF4G1 (residues 682-1599), with various amounts of WT-eIF4B or M1-eIF4B (0-0.5 μM), in buffer containing 20 mM tris-acetate pH 7.5, 100 mM potassium acetate, 2 mM Mg acetate, 10% glycerol, 2 mM ATP-Mg^2+^, and 1 mM DTT. The results shown are averages of three independent experiments with SEM.

### Fluorescence Anisotropy Assay

The anisotropy binding assay was done as described previously.^40,43^ The reaction contained 10 nM labeled WT- or M1-eIF4B with 0–125 nM 40S subunit, or 20 nM labeled RNA with 0–2000 nM WT- or M1-eIF4B, in buffer containing 20 mM tris-acetate pH 7.5, 70 mM KCl, 2 mM MgCl_2_, 0.1 mM spermidine, 0.1 mg/ml BSA, 10% glycerol, and 1 mM DTT. Fitting with a quadratic equation suggests a tight dissociation constant below the lower limit (< 10 nM) for both eIF4Bs. The results shown are averages of three independent experiments with SEM.

## Supplementary Figure Titles and Legends

**Supplementary Figure 1.**
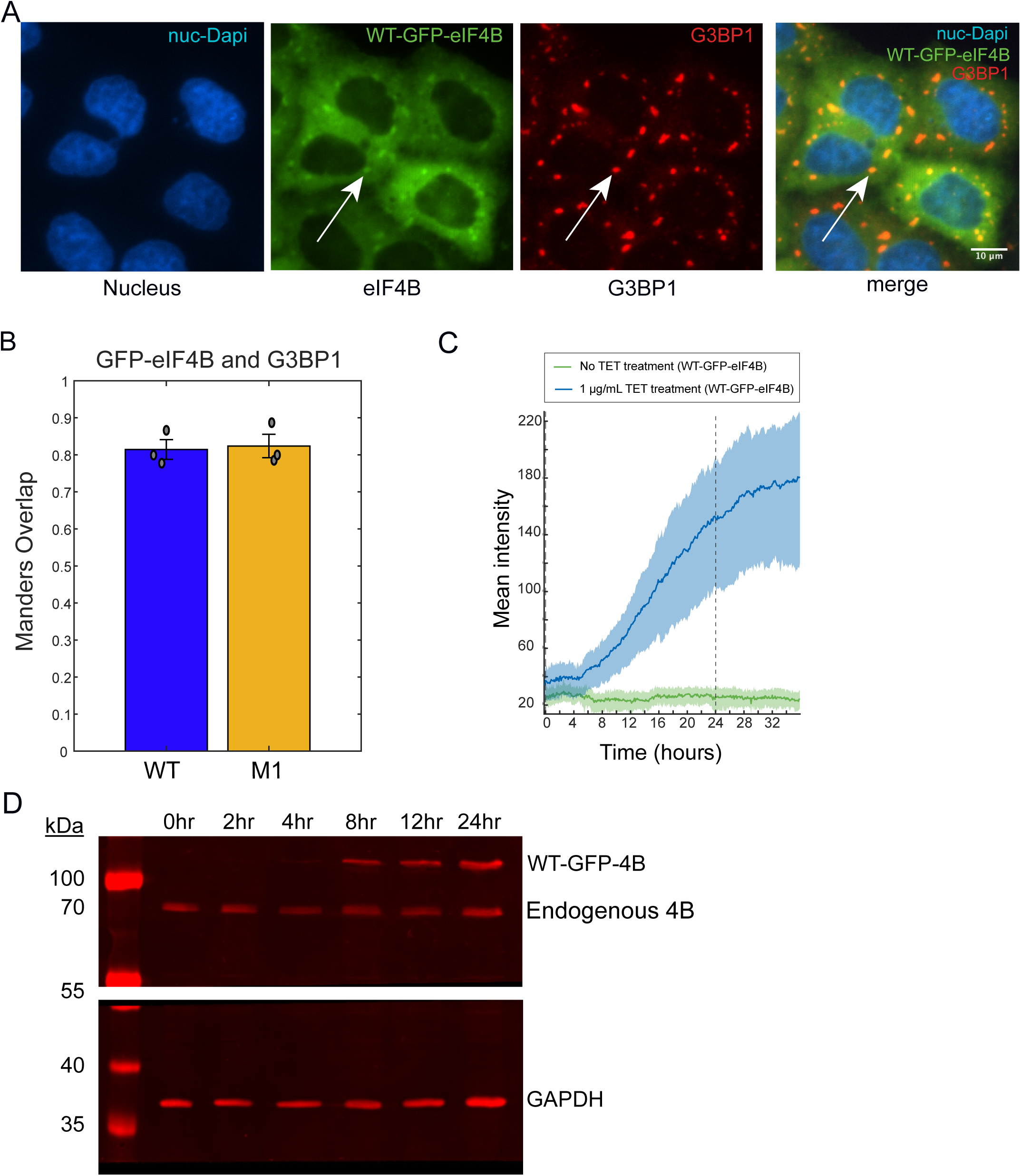
Characterizing eIF4B overexpression cell line. (A) WT-GFP-eIF4B cells were subjected to immunofluorescence (IF) to visualize the stress granule nucleator G3BP1 using an anti-G3BP1 antibody, in order to verify GFP-eIF4B localization in stress granules. The images, from left to right, are as follows: DAPI (nucleus), WT-GFP-eIF4B, G3BP1 (SG marker), and the merged image (both eIF4B and G3BP1). (B) Immunofluorescence data were analyzed using Mander’s overlap of WT-GFP-eIF4B (blue) and M1-GFP-eIF4B (orange) granule signals with G3BP1 (SG marker) after cells were treated with 125 μM sodium arsenite for 2 hours. The mean and SEM of 3 replicates are shown; gray dots represent each replicate. (C) Measuring mean intensity of WT-GFP-eIF4B TReX Flip-In tetracycline inducible cells over a 36 hour time frame. Dark lines represent the mean and the shaded regions are 25th/75th quartiles of data for cells in the respective condition. All subsequent experiments used a 24-hour induction time to express GFP-eIF4B, which is indicated at the dotted line. (D) Western blot was performed at different induction times to measure the presence of endogenous eIF4B and overexpressed WT-GFP-eIF4B, as indicated. GAPDH is used as a loading control. The gel was cut in two so that the primary antiserum to eIF4B (mouse) and GAPDH (goat) could be used. From left to right is the time of induction 0–24 hours. 30 μg of total protein was loaded per well and separated using a 8% SDS page gel.

**Supplementary Figure 2.**
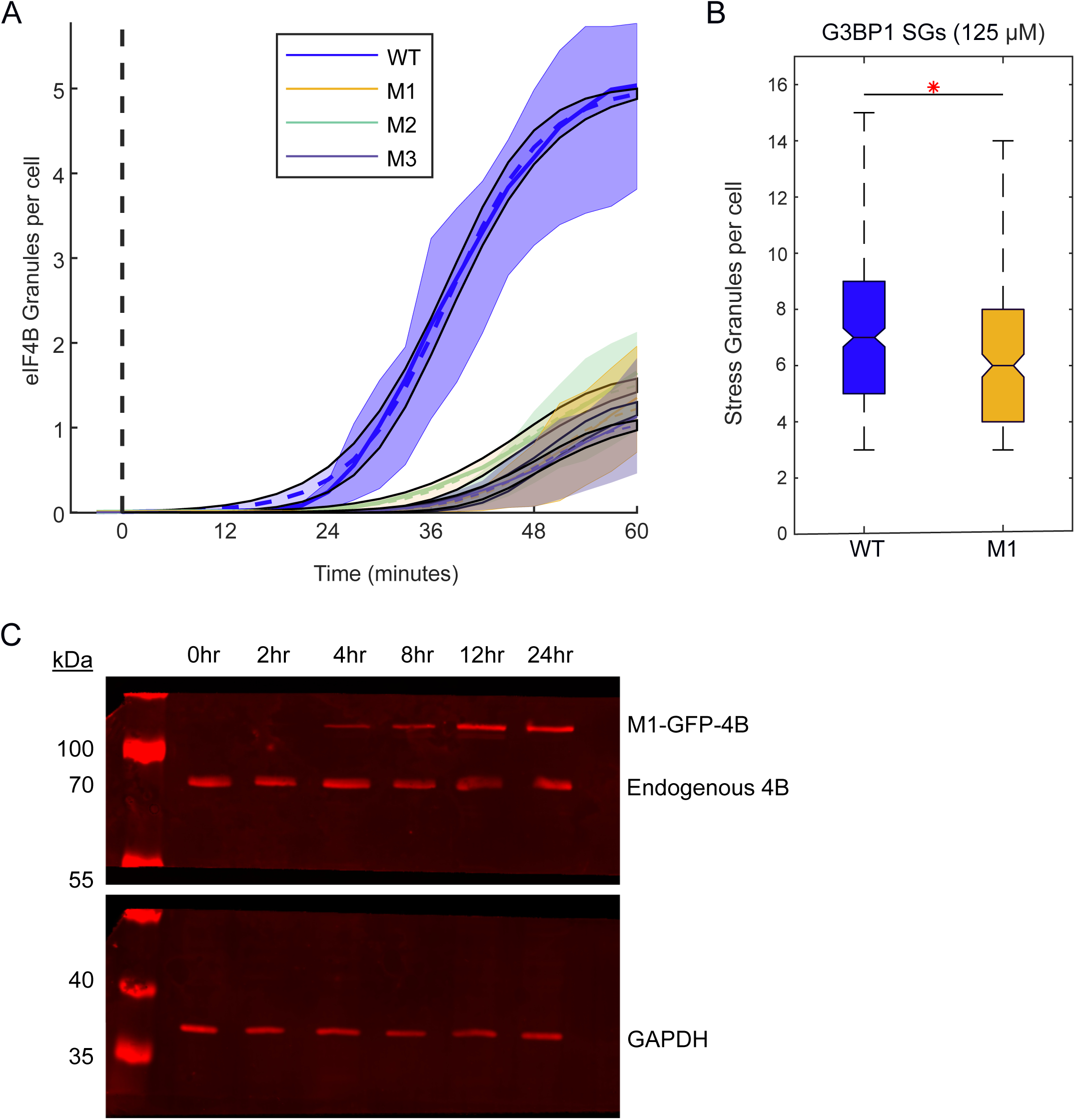
Characterizing the RRM of eIF4B overexpression cell line. (A) SG formation equations were fit to measure the f max as each cell line approaches 1 hour. Mean (dark line) and 25th/75th quartiles (shaded regions) of SGs formed per cell line over time following treatment with 125 μM sodium arsenite. WT-GFP-eIF4B in red, M1-GFP-eIF4B (F139A) in green, M2-GFP-eIF4B (F99A/N102A/R135A) in yellow, and M3-GFP-eIF4B (R135A/K137A/F139A) in blue. Sodium arsenite treatment addition time point shown by dashed line (hour 0). The number of SGs per cell for each cell line WT=5.04, M1 =1.35, M2 =1.65, and M3 =1.13. (B) Box and whiskers plot showing G3BP1 SGs in the WT and M1 cell lines following 2 hours of treatment with 125 μM arsenite. Black line with the red asterisk indicates a significant difference between the groups (P = 0.0016). Significance was determined via One-Way ANOVA comparison. (C) Western blot was performed at different induction times to measure the presence of endogenous eIF4B and overexpressed M1-GFP-eIF4B, as indicated. GAPDH is used as a loading control. The gel was cut in two so that the primary antiserum to eIF4B (mouse) and GAPDH (goat) could be used. From left to right is the time of induction 0–24 hours. 30 ug of total protein was loaded per well and separated using a 8% SDS page gel.

**Supplementary Figure 3.**
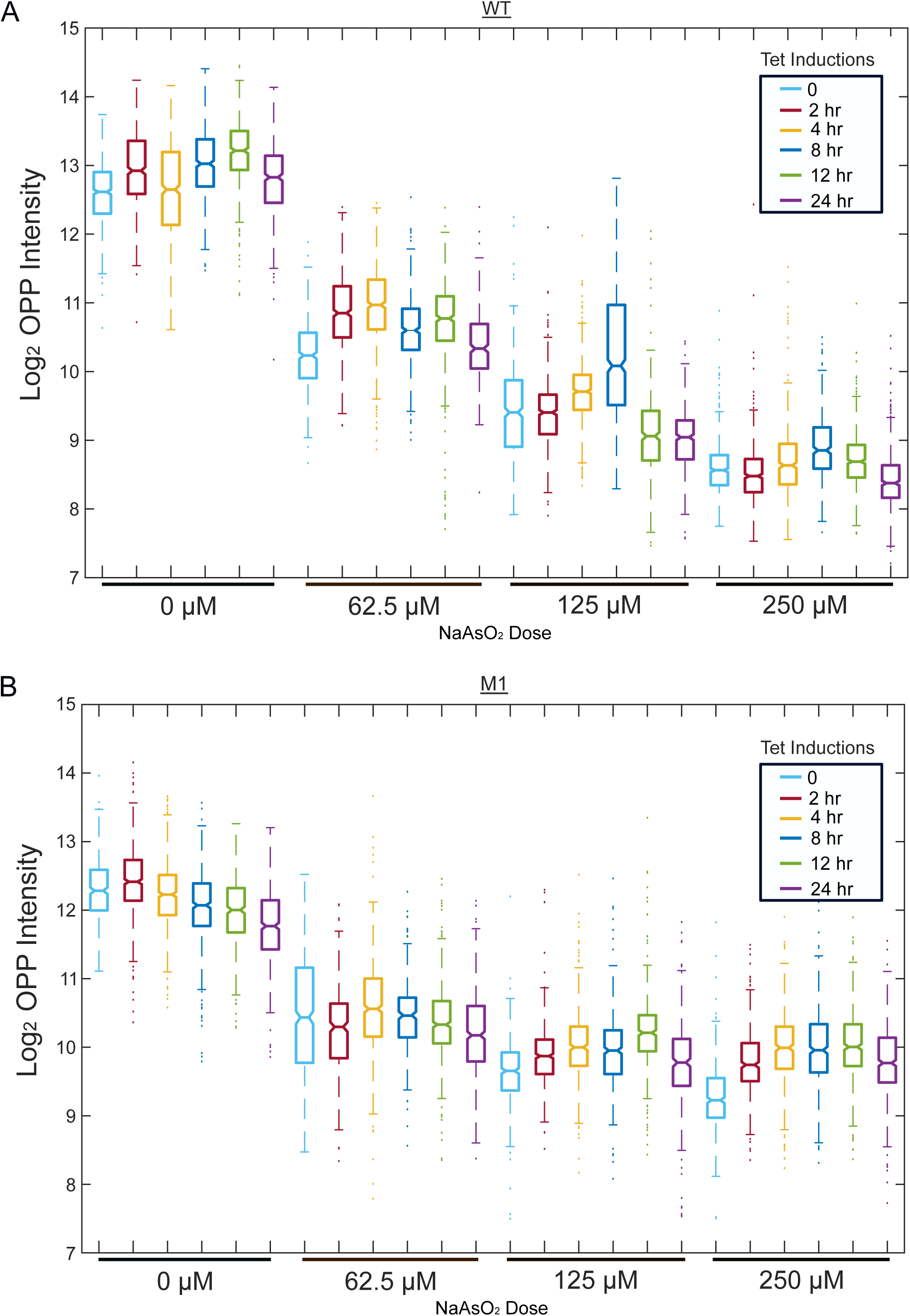
Measuring translation activity in WT and M1 cells. (A) Box and Whisker plot of single-cell Log_2_ O-propargyl-puromycin (OPP) signal intensity of WT-GFP-eIF4B (WT) HeLa cells at varying TET inductions (In colors ranging from blue to purple) and grouped by sodium arsenite doses as indicated (62.5–250 μM). Colors indicate hours of TET induction with 0 hours (light blue), 2 hours (red), 4 hours (dark blue), 12 hours (green), and 24 hours (purple). Measurements were made after one hour and 30 mins of sodium arsenite treatment and OPP was added to the last 30 minutes of the experiment (total experiment of 2 hours). (B) Box and Whisker plot of single-cell Log_2_ O-propargyl-puromycin (OPP) signal intensity of M1-GFP-eIF4B (M1) HeLa cells at varying TET inductions (In colors ranging from blue to purple) and grouped by sodium arsenite doses as indicated (62.5–250 μM). Colors indicate hours of TET induction with 0 hours (light blue), 2 hours (red), 4 hours (dark blue), 12 hours (green), and 24 hours (purple). Measurements were made after one hour and 30 mins of sodium arsenite treatment and OPP was added to the last 30 minutes of the experiment (total experiment of 2 hours).

**Supplementary Figure 4.**
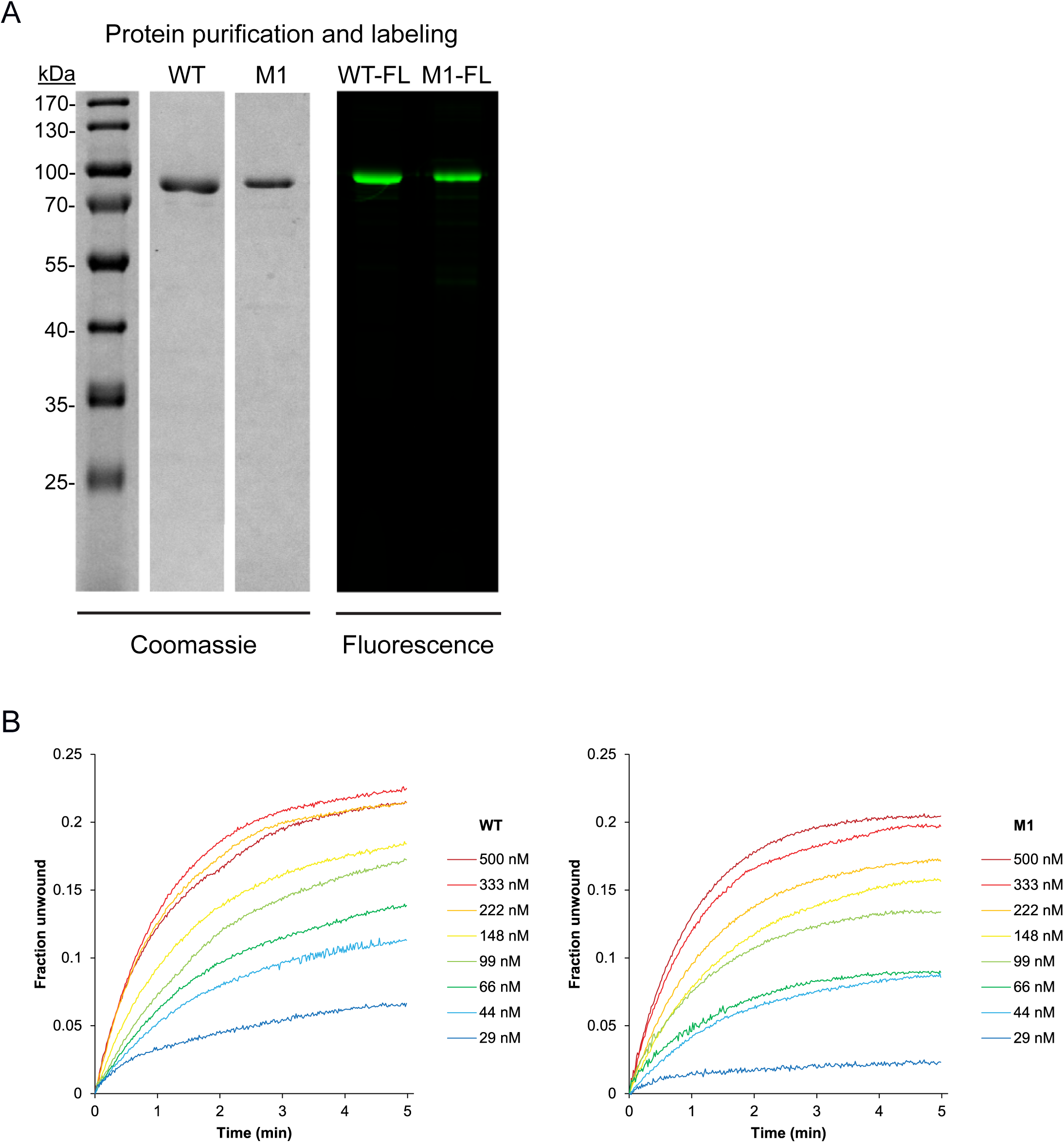
Protein purification and Labeling. (A) Coomassie-stained SDS-PAGE of purified WT-eIF4B and M1-eIF4B (left), and fluorescence imaging of a gel showing fluorescently labeled WT-eIF4B and M1-eIF4B (right). These protein preparations were used for the experiments shown in Figure 4. (B) Real-time fluorescent helicase assay where the ATP-dependent unwinding of a synthetic RNA duplex is monitored in the presence of purified eIF4A helicase, eIF4G, and WT-eIF4B or M1-eIF4B as indicated.

